# Sub-lethal aquatic doses of pyriproxyfen may increase pyrethroid resistance in malaria mosquitoes

**DOI:** 10.1101/2021.01.11.426299

**Authors:** Mercy A. Opiyo, Halfan S. Ngowo, Salum A. Mapua, Monica Mpingwa, Nancy S. Matowo, Silas Majambere, Fredros O. Okumu

## Abstract

**Background:** Pyriproxyfen (PPF), an insect growth hormone mimic is widely used as a larvicide and in some second-generation bed nets, where it is combined with pyrethroids to improve impact. It has also been evaluated as a candidate for auto-dissemination by adult mosquitoes to control *Aedes* and *Anopheles* species. We examined whether PPF added to larval habitats of pyrethroid-resistant malaria vectors can modulate levels of resistance among emergent adult mosquitoes.

**Methodology:** Third-instar larvae of pyrethroid-resistant *Anopheles arabiensis* (both laboratory-reared and field-collected) were reared in different PPF concentrations, between 1×10^-9^ milligrams active ingredient per litre of water (mgAI/L) and 1×10^-4^ mgAI/L, or no PPF at all. Emergent adults escaping these sub-lethal exposures were tested using WHO-standard susceptibility assays on pyrethroids (0.75% permethrin and 0.05% deltamethrin), carbamates (0.1% bendiocarb) and organochlorides (4% DDT). Biochemical basis of pyrethroid resistance was investigated by pre-exposure to 4% PBO. Bio-efficacies of long-lasting insecticide-treated nets, Olyset^®^ and PermaNet 2.0 were also examined against adult mosquitoes with or without previous aquatic exposure to PPF.

**Results:** Addition of sub-lethal doses of PPF to larval habitats of pyrethroid-resistant *An. arabiensis*, consistently resulted in significantly reduced mortalities of emergent adults when exposed to pyrethroids, but not to bendiocarb or DDT. Mortality rates after exposure to Olyset^®^ nets, but not PermaNet 2.0 were also reduced following aquatic exposures to PPF. Pre-exposure to PBO followed by permethrin or deltamethrin resulted in significant increases in mortality, compared to either insecticide alone.

**Conclusions:** Partially-resistant mosquitoes exposed to sub-lethal aquatic concentrations of PPF may become more resistant to pyrethroids than they already are without such pre-exposures. Studies should be conducted to examine whether field applications of PPF, either by larviciding or other means actually excercebates pyrethroid-resistance in areas where signs of such resistance already exist in wild the vector populations. The studies should also investigate mechanisms underlying such magnification of resistance, and how this may impact the potential of PPF-based interventions in areas with pyrethroid resistance.

## Introduction

Insecticides remain an important asset for malaria control programs. In countries where malaria is largely endemic in Africa, insecticide based interventions primarily insecticide-treated nets (ITNs) or now long-lasting insecticide-treated nets (LLINs) and indoor residual spraying (IRS) are the most widely used vector control tools [1]. Both the LLINs and IRS have been and they remain in the frontline in the fight against malaria and they are associated with a significant reduction in the malaria burden observed between 2000 and 2015 [2, 3]. However, in recent years, malaria burden has been on the rise again and further gains have slowed down [4, 5]. Therefore, there is an urgent need to investigate why we are observing these changes from different intervention dimensions to design strategic measures to recover the deteriorating impact of the frontline interventions.

According to the World Health Organization Pesticide Evaluation Scheme (WHOPES), the second primary intervention, IRS can be done with four classes of insecticides (pyrethroids, carbamates, organophosphates, and neonicotinoids) [6]. For bed nets, pyrethroids remain the key class of insecticide approved for use to date, although in recent years, there has been significant progress in search of more active ingredients for combination with pyrethroids in bed nets. Notable examples include pyrethroid-combination based bed nets such as Royal Guard^®^ (alpha-cypermethrin and pyriproxyfen), Olyset^®^ Plus (permethrin and Piperonyl Butoxide (PBO)), PermaNet^®^3.0 (deltamethrin and PBO), and Interceptor ^®^ G2 (alpha-cypermethrin and Chlorfenapyr) nets have been proposed [6]. However, excessive reliance on pyrethroids as the most common active ingredient in both public health and agricultural sectors has created major selection pressures and has led to the rapid expansion of pyrethroid resistance among malaria vector populations [7–10]. And to date, pyrethroid resistance in malaria vector populations has been documented nearly everywhere across all malarious settings in Africa [7]. This poses a challenge to malaria control efforts made so far and urgent strategic measures are needed. Even though intense and continuous efforts in search of more, new, safe, and effective insecticidal active ingredients are under investigation, this will take time before they are ready and are fully commercialized [6].

Insecticide resistance management strategies commonly promote the use of two or more insecticide classes, either as combinations, mosaics, or mixtures, to increase the responsiveness of target vectors to insecticide-based interventions [7]. Using two different insecticides with different modes of action can delay the development of resistance as well as delay the spread of resistant alleles in the vector populations. Other promising strategies to combat or delay insecticide resistance spread includes the use of synergists such as Piperonyl-Butoxide (PBO), an inhibitor of Cytochrome P450 monooxygenases, and commonly associated with phenotypic pyrethroid resistance [11, 12]. For example, Olyset^®^ Plus is a new generation long-lasting insecticide-treated net (LLIN), made of polyethylene netting incorporating permethrin and PBO as a synergist, and has been shown to have higher efficacy than regular permethrin-based LLINs against malaria mosquitoes bearing metabolic resistance to pyrethroids [13, 14]. Other alternative pyrethroid-based insecticide resistance mitigating LLINs such as Interceptor^®^ G2 (pyrethroid-Chlorfenapyr) and Royal Guard^®^ (alpha-cypermethrin and pyriproxyfen) have been evaluated [15], and more studies are underway to assess their impacts. Unfortunately, the cost of these nets compared to the standard LLIN’s [16] still limits there use. Another alternative strategy, also widely utilized in the past, is larval source management. This involves reducing the vector population density by killing or preventing immature stages of mosquitoes while in their aquatic habitats from becoming adults, by the use of biological agents and chemicals [17–19]. While this strategy has been used with great success in some settings including Africa, it is faced by logistical challenges, such as identifying the diverse and dynamic Anopheline larval sites as well as the cost incurred for delivering the larvicides [17, 20]. More so, its inclusion in malaria control programmes has been recently a topical agenda in most malaria vector control decision-making platforms.

Pyriproxyfen (PPF) is a juvenile insect growth inhibitor hormone mimic, which prevents embryo development and also stops metamorphosis in insects [21]. It has been certified by the Word Health Organization (WHO) for larval control due to its high effectiveness in very small doses, and its safety to humans [22]. PPF reduces both longevity and fecundity of mosquitoes but also causes sterilization in adult females of *Anopheles gambiae* [23, 24], *An. arabiensis* [25, 26] and non-malaria mosquitoes such as *Culex quinquifasciatus* [27] and *Aedes albopictus* [28]. This juvicidal insecticide can also be combined with other insecticide classes such as pyrethroids to reduce further spread of resistant alleles in mosquito populations, by reducing fecundity and longevity, sterilizing pyrethroid-resistant female strains that escape the effect of pyrethroid insecticide, thereby reducing the next generation of vectors. This strategy, where the target insect is highly resistant to one compound but susceptible to another in an insecticide mixture, has been tested in adult malaria vectors resistant for pyrethroids with great success [29]. Pyriproxyfen has also been used previously in combination with pyrethroids on bed nets to target insecticide-resistant malaria vectors. An example is Olyset^®^ Duo, which combines permethrin and pyriproxyfen, and has been shown to achieve higher mortalities on pyrethroid-resistant *Anopheles* than the regular Olyset^®^ Net with just permethrin alone [23, 24, 30, 31]. Studies on these new generations of bed nets however still need further evaluation, before the nets can be fully rolled out for malaria control [32].

PPF can also be used as a larvicide, either by direct application [33] or auto-dissemination by female mosquitoes into larval habitats [34, 35]. Studies have shown that it is possible to contaminate adult mosquito resting sites, such that these mosquitoes are able to carry small but effective doses of PPF on their legs, and subsequently deliver these into the larval sites, during egg-laying, a strategy often known as auto-dissemination [35–37]. This technology has been tested in a small field trial in Peru with great success [34], and also in Brazil against vectors of dengue and Zika virus [38] as well as in Tanzania against malaria mosquitoes (Opiyo et al., unpublished data). It has also been demonstrated that adult mosquitoes can ingest PPF when combined with attractive sugar baits (ASB) and feacally disseminate into water consequently reducing adult emergence [39].

Given the growing interest in PPF as a potential complementary malaria control chemical, it is vital to evaluate the impact of combining PPF with already existing pyrethroid based tools, and how such combinations may interact, especially in areas with insecticide resistance. For example, what would happen if PPF were used in the same communities where LLINs or other pyrethroid-based interventions are already widely used? How would the mosquitoes exposed to PPF in their aquatic stages respond to LLINs and/or IRS? What would be the optimal combination of chemicals used as larvicides and adulticides be? To begin addressing these questions, we explored whether PPF added to larval habitats of pyrethroid-resistant malaria vectors would reduce or increase the level of resistance among adult mosquitoes emerging from these habitats and whether common LLINs would be more or less effective to mosquitoes previously exposed to PPF. Our findings are expected to help guide selections of interventions to mitigate resistance and integrated vector management approach against mosquito-borne diseases.

## Materials and Methods

### Study location

This study was conducted at Ifakara Health Institute (IHI), inside the Vector Biology and Control facility, the *VectorSphere*.

### Mosquito larvae

Wild resistant *An. arabiensis* larvae were sampled weekly from Namwawala village located in the flood plains of the Kilombero River (8.1°S and 36.6°E) in south-eastern Tanzania. The village experiences a long rainy season between February and May with a short rainy season between November and December. Polymerase Chain Reaction (PCR) assays on *Anopheles gambiae* s.l collected from this village has consistently returned as 100% *An. arabiensis* for the past 9 years, so all *An. gambiae* adults obtained consisted *An. arabiensis*. Communities living here are subsistence farmers growing mainly maize and rice.

Larvae of mixed ages were collected from both rice fields and breeding pools located outside the rice farms between June and December 2016. Larvae were transported to the *VectorSphere* for sorting and rearing. In the *VectorSphere*, third-instar larvae were collected immediately upon arrival and used in the subsequent experiment, and the remaining young larvae were reared to third instar larvae under 27°C±2°C until ready for use. A sub-sample of the field samples were reared to adults and also used in the subsequent experiments. Third-instar larvae of pyrethroid-resistant *An. arabiensis* was also obtained from the IHI mosquito colony (Kining’ina colony or Mosquito City colony). This *An. arabiensis* colony was established in 2007 but was found to be highly resistant to pyrethroids in 2015 (Matowo et al., unpublished). The basis of this resistance is assumed to be biochemical and mediated by elevated monooxygenases, as it is reversible with the synergist, Piperonyl Butoxide (PBO). The Kining’ina (Mosquito City) colony strain is reared in a semi-field system (SFS) condition, under natural temperature and light-dark cycles. The humidity for the Kining’ina colony was artificially increased to 70-80% for the adult mosquitoes. The colony is routinely reared on 10% glucose, and blood meal for colony maintenance is provided by a human arm daily, and the larvae fed on Tetramin^®^ fish food twice a day.

### Treatments with pyriproxyfen (PPF)

The PPF used in this study is a dust formulation containing 10 % active ingredient (AI). Before the actual experiment, and to determine the range of sub-lethal concentrations to work with, we first initially reared 3^rd^ instar larvae from the *An. arabiensis* colony (3 replicates), as well as field samples (25 larvae for 5 replicates), to a wide range of PPF concentrations and controls. We aimed to select sub-lethal doses with minimum effect on adult emergence so that we would have enough adult females to test. Concentrations between 1×10^-7^ mg AI per litre (mgAI/L) and 1×10^-2^mgAI/L were tested initially. However, during the actual experiments, we also included 1×10^-8^ mgAI/L and 1×10^-9^ mgAI/L. In the entire actual experiment, not all concentrations were used for rearing larvae due to lack of sufficient larvae. All test concentrations were prepared from serial dilutions of standard stock solutions, freshly made for each replicate as follows: we prepared stock solution by adding 1mg of 10% AI PPF in 1 litre of water and left the stock solution on the shaker for 24 hours to thoroughly mix giving a stock solution of 0.1 mgAI/L. After determining the emergence of larvae in different doses, we chose the dose ranges that yielded at least 20% emergence (i.e. dose ranges from 1×10^-4^ mgAI/L to 1×10^-9^ mgAI/L) for subsequent experiments.

For the WHO-susceptibility tests, 3rd instar larvae obtained from the *An. arabiensis* laboratory colony, as well as from the field were therefore reared in pools containing 1200 larvae in different PPF doses, ranging from 1×10^-4^ mg AI/L to 1×10^-9^ mgAI/L and effects monitored. During this period, the larvae were fed on Tetramin^®^ fish food, as was normal practice in the insectary colony. The adult mosquitoes emerging from basins with these different PPF treatments were kept separately and provided with 10% glucose. The adults were later subjected to WHO-standard susceptibility assays to determine if they were more or less resistant to the same concentrations of the insecticides. In all tests, we also maintained similar quantities of larvae in pools without any PPF treatment, to act as controls.

### WHO insecticide susceptibility assays

Three to five days old non-blood fed female mosquitoes emerging from PPF treatments (dose range in decreasing order: 1×10^-4^ mg AI/L, 1×10^-5^ mg AI/L, 1×10^-6^mg AI/L, 1×10^-7^mg AI/L and 1×10^-8^mg AI/L) were tested following WHO standard susceptibility test protocol [40], against three insecticide classes namely: Type I and II pyrethroids (0.75% permethrin and 0.05% deltamethrin respectively), Organochlorine (4% DDT), and Carbamate (0.1% bendiocarb). Parallel control assays were run using non-treated papers alongside all insecticides tested. For each PPF dose and the controls, one hundred emergent mosquitoes, in batches of 25 mosquitoes each (i.e. 4 replicates) were exposed to each of the candidate insecticides. Control assays, using non-insecticidal test papers as recommended in the WHO protocols, were run in parallel each time with the same number of mosquitoes. To quantify and determine the effects of PPF treatment in emergent adults, we also performed susceptibility assays on adults emerging from non-PPF treated larval habitats using the procedures already described. All tests on the laboratory-reared mosquitoes and field mosquitoes were done using similar WHO susceptibility assay protocols inside the *VectorSphere*.

### LLIN bio-efficacy assays

We also evaluated the bio-efficacy of WHO-recommended Long-Lasting Insecticide Treated Nets (LLINs), i.e. Olyset^®^ Net (active ingredient is permethrin incorporated into polyethylene fibres at 2% w/w; Sumitomo Chemical Co., Ltd., Osaka, Japan). A non-insecticidal net type, Safi^®^ Net (A to Z, Arusha, Tanzania) was used as a control. All the bed nets used for this assay were new (not previously used) and had been purchased from local stores. The tests were conducted using standard WHO recommended plastic cones [41]. Similar to the WHO susceptibility assays, we used female mosquitoes emerging from different PPF-treatments (dose ranges 1×10^-8^ mg AI/L to 1×10^-5^ mg AI/L). Batches of five female *An. arabiensis* mosquitoes (three to five days old and not previously blood-fed) emerging from larval basins with different PPF concentrations were exposed for 3 minutes to each of the 5 sides of each LLIN type. For each PPF treatment, we replicated the mosquito exposures at least ten times, each time using a new batch of mosquitoes (each side n=50 mosquitoes, each net type ranges; n=200-250 mosquitoes). The cone assays were run at 27 ± 1°C and 70 ± 10%. Knockdown was recorded after 60 minutes, and mosquito mortality, recorded after 24 hours’ post-exposure. To quantify the effects of PPF treatment on the efficacy of the LLINs, we also conducted similar bioassays on the same nets as described above using female mosquitoes without prior aquatic exposure to PPF treatments.

### Tests using the synergist, Piperonyl Butoxide (PBO), to assess the biochemical basis of resistance

To assess whether the observed pyrethroid resistance was associated with any underlying elevation of enzymes metabolizing the pyrethroids and whether effects of PPF can be reversed by adding synergists; we performed additional synergism assays with PBO, an inhibitor of cytochrome P450 detoxifying enzymes and esterases [42, 43]. *An. arabiensis* collected as 3^rd^ instar larvae from both the field (Namwawala village) and laboratory colony were reared to adults as described above. Four sets of standard WHO susceptibility tests were performed by exposing mosquitoes to 0.05% deltamethrin and 0.75 % permethrin test papers alone. Another two sets of tests were conducted by first pre-exposing the mosquitoes for 1 hour to 4% PBO, followed by another 60 minutes’ exposure to either 0.75% permethrin or 0.05% deltamethrin. There were three batches of control mosquitoes as follows: a batch of mosquitoes exposed to plain white papers to act as environmental controls, to monitor any environmental contamination, a second batch exposed to papers impregnated with olive oil, to control for the insecticidal content of the test papers, and the third batch of mosquitoes exposed to papers treated with 4% PBO alone, to control for effects of the synergist on its own. Mosquito knockdown rates were recorded every 5 minutes for up to 1 hour during exposure, and mortality recorded 24 hrs after exposure. During the 24 hrs period, the mosquitoes were provided with 10% glucose solution. The synergist test was done with pyrethroids alone and not any of the other insecticide classes.

### Data analysis

The results were analyzed using open source software, R version 3.3.2 [44]. First, data from dose-response assays were pooled by PPF concentrations, for each mosquito source (i.e. *An. arabiensis* colony, and *An. arabiensis* field mosquitoes), and adult emergence rates taken as the primary outcome. Susceptibility bioassay data for mosquito populations pre-exposed to PPF of different concentrations and those without pre-exposure to PPF were first summarised as mean percentage (%) mortality monitored 24-h post-exposure per mosquito source and insecticide type. Thereafter, the mosquito populations emerging from different PPF concentrations were pooled per insecticide and mosquito source and the mean % mortality observed 24-h post-exposure and comparison made between mosquito populations pre-exposed to PPF and those without pre-exposure to PPF using paired sample t-test. All the data were summarized as percentage mean emergence. For WHO-susceptibility and bio-efficacy (cone) assay [40], test results were analyzed as 24-hr percentage mean mortality as described in WHO-susceptibility test procedures, with mortality as the primary outcome. For synergist tests, the time taken to knock-down 50 % (KD50) of the mosquito population was estimated using log-probit analysis at 95% CI and 24 hr mortality evaluated as described above.

### Ethical approval

This study was approved by the Ifakara Health Institute’s Review Board (IHI-RB) (IHRDC/IRB/NO. A-32 & IHI/IRB/No: 34-2014) and the Medical Research Coordinating Council (MRCC), at the National Institute of Medical Research (NIMR) in Tanzania (NIMR/HQ/R.8a/Vol. IX/764 & NIMR/HQ/R.8a/Vol.IX/1903).

### Consent for publication

Permission to publish this work was also obtained from the National Institute of Medical Research (NIMR), REF: NIMR/HQ/P.12 VOL XXXI/57.

## Results

### Initial tests to determine the range of sub-lethal PPF doses

Emergence rates of third-instar larvae of field-collected *An. arabiensis*, and laboratory-reared *An. arabiensis* exposed to a range of PPF doses are summarized in Fig.1. Data from five replicates of field *An. arabiensis* and three replicates of *An. arabiensis* colony was pooled per dose to estimate the dose that would allow at least 5 % emergence to be used in the subsequent experiments. Concentrations of 1×10^-2^ mg AI/L and 1×10^-3^ mg AI/L completely inhibited the emergence of adult mosquitoes from treated pools, in tests with the laboratory-reared and field-collected mosquitoes (Fig.1). However, the PPF doses between 1×10^-7^ mg AI/L and 1×10^-4^ mg AI/L allowed 5 % to nearly 50 % emergence. All subsequent experiments after this initial test therefore used doses in the range of 1×10^-8^ mg AI/L and 1×10^-4^ mg AI/L.

**Fig. 1:**
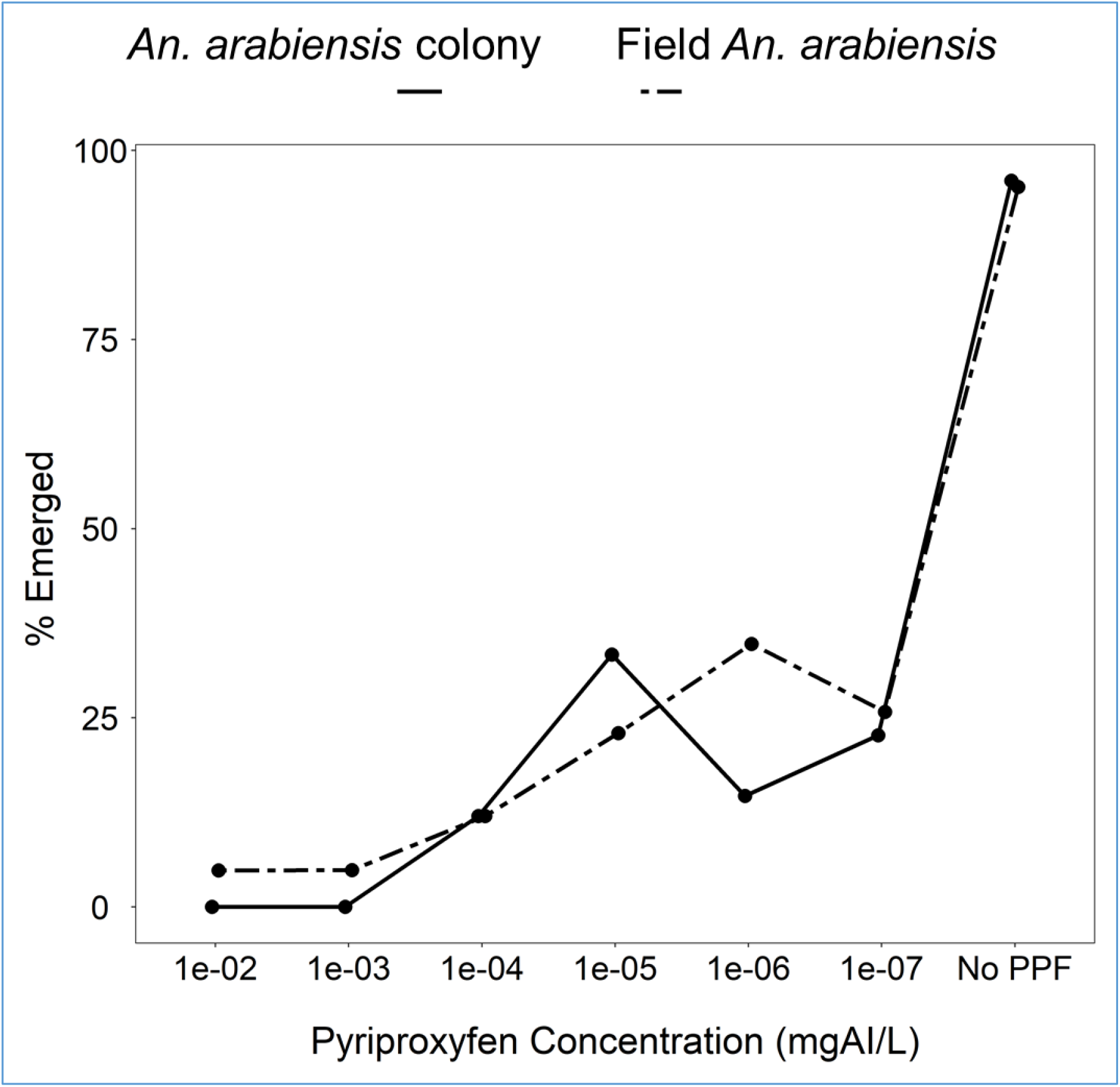
Percentage of mosquitoes emerging as adults from larval-rearing basins treated with different concentrations of pyriproxyfen (PPF): *Anopheles arabiensis*, from Kining’ina colony population; and field *Anopheles arabiensis*, from Namwawala village.

### WHO insecticide susceptibility bio-assays

Results of the standard WHO-susceptibility tests using mosquitoes emerging from PPF-treated and untreated basins are summarized in (Figs. 2, 3 & 4). Susceptibility to deltamethrin, permethrin, DDT, and bendiocarb was assessed for *An. arabiensis* colony, and also for field-collected *An. arabiensis* mosquitoes (Figs. 2, 3 & 4). Without any exposure to PPF, the 24-hr mortality of the laboratory-reared *An. arabiensis* colony was 35% and 83% in tests using permethrin and deltamethrin treated papers respectively confirming the pyrethroid resistance in these colonies (Figs. 2 & 3). The same mosquitoes were however susceptible to DDT (24-hr mortality = 98%) and bendiocarb (24-hr mortality = 98 %) (Fig. 3). Similarly, without any pre-exposure to PPF, the 24-hr mortality of the field-collected *An. arabiensis* mosquitoes were susceptible to only bendiocarb (98 %) (Fig. 4). These field-collected mosquitoes were resistant to permethrin (24-hr mortality = 27%), deltamethrin (24-hr mortality = 82%), and DDT (24-hr mortality = 45 %) (Figs. 2 & 4). When the *An. arabiensis* colony pre-exposed to PPF were pooled for all concentrations and compared with those without pre-exposue to PPF, there was significant difference in 24-h mortality following exposure to deltamethrin (t=4.012, df=5.412, p=0.0086), while there was a reduction in 24-h mortality when exposed to permethrin but the differences were not statiscally significant (t=1.710, df=3.288, p=0.178) (Fig.2). On the ther hand, when the field *An. arabiensis* population pre-exposed to PPF were pooled for all concentrations and compared with those without pre-exposure to PPF, there was significant difference in 24-h mortality following exposure to deltamethrin (t=3.693, df=6.367, p=0.0091), while when exposed to permethrin, there was a reduction in 24-h mortality but this difference was not statistically significant (t=1.069, df=5.487, p=0.333) (Fig.2).

**Fig. 2:**
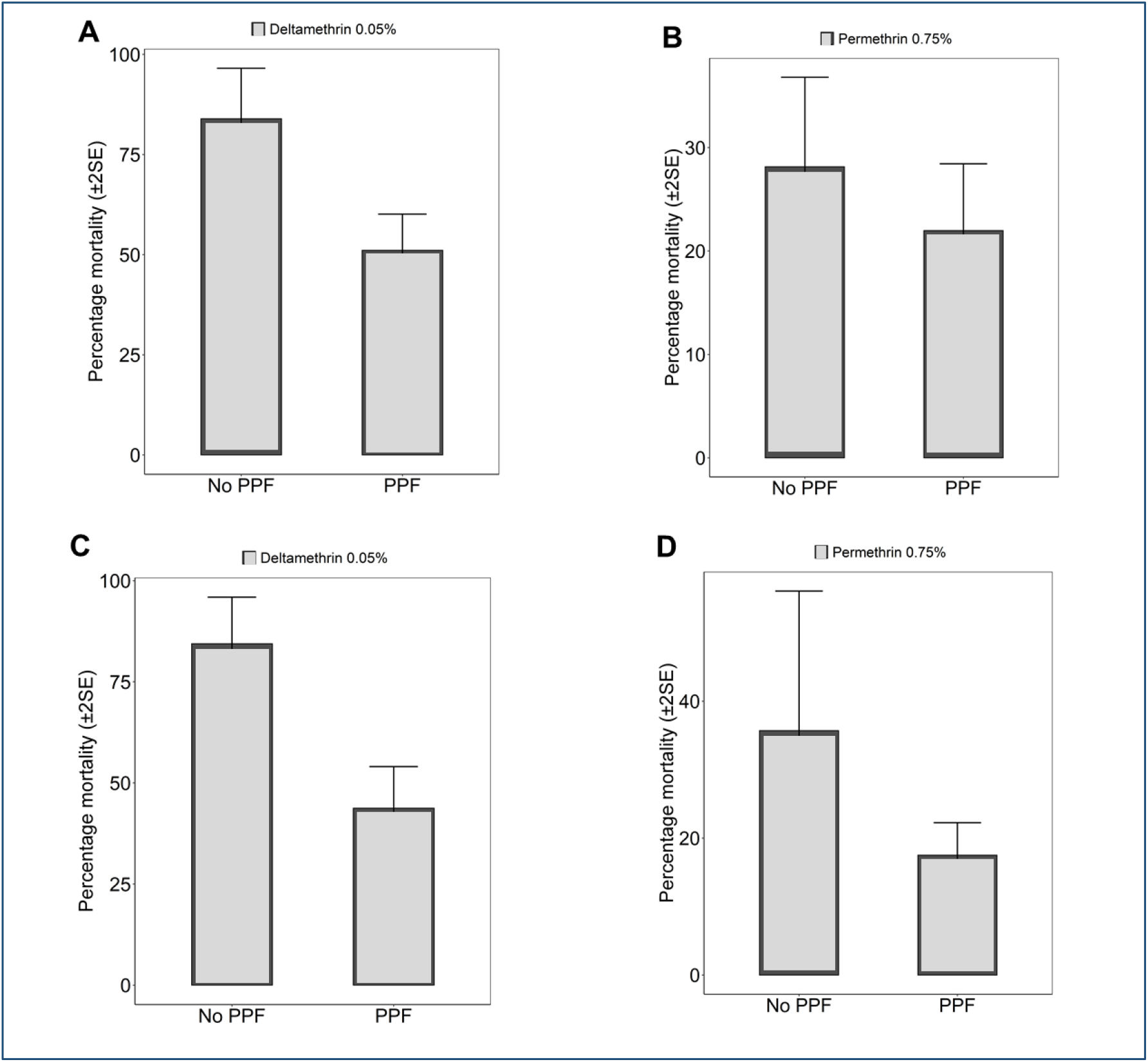
Pooled percentage mortality (24-hour) of: **A & B**. field resistant *An. arabiensis* mosquitoes, and **C & D.** resistant *An. arabiensis* colony reared through different doses of PPF (mg of active ingredient per litre, i.e. mgAI/L) and exposed to: **0.75% Permethrin**, **0.05% Deltamethrin**, using standard WHO susceptibility testing guidelines. Bars labelled “No PPF” represent percentage mortality of mosquitoes from the same colony, which were not treated with PPF but were exposed to the same insecticide at standard WHO-recommended dose.

**Fig. 3:**
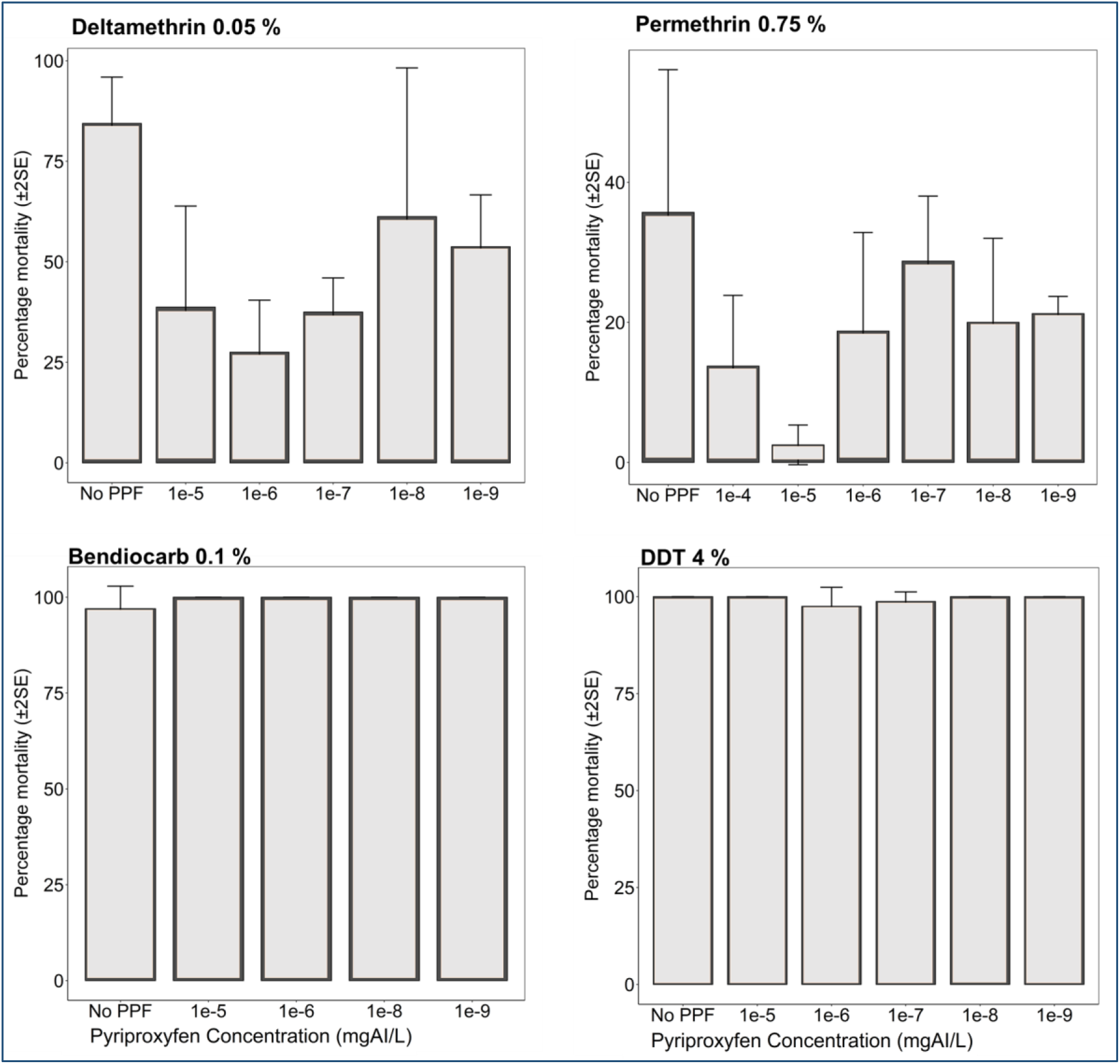
Percentage mortality (24-hour) of resistant *An. arabiensis* colony mosquitoes reared through different doses of PPF (mg of active ingredient per litre, i.e. mgAI/L) and exposed to commonly used insecticides using standard WHO susceptibility testing guidelines. Bars labelled “No PPF” represent percentage mortality of mosquitoes from the same colony, which were not treated with PPF but were exposed to the same insecticide at standard WHO-recommended dose.

**Fig. 4:**
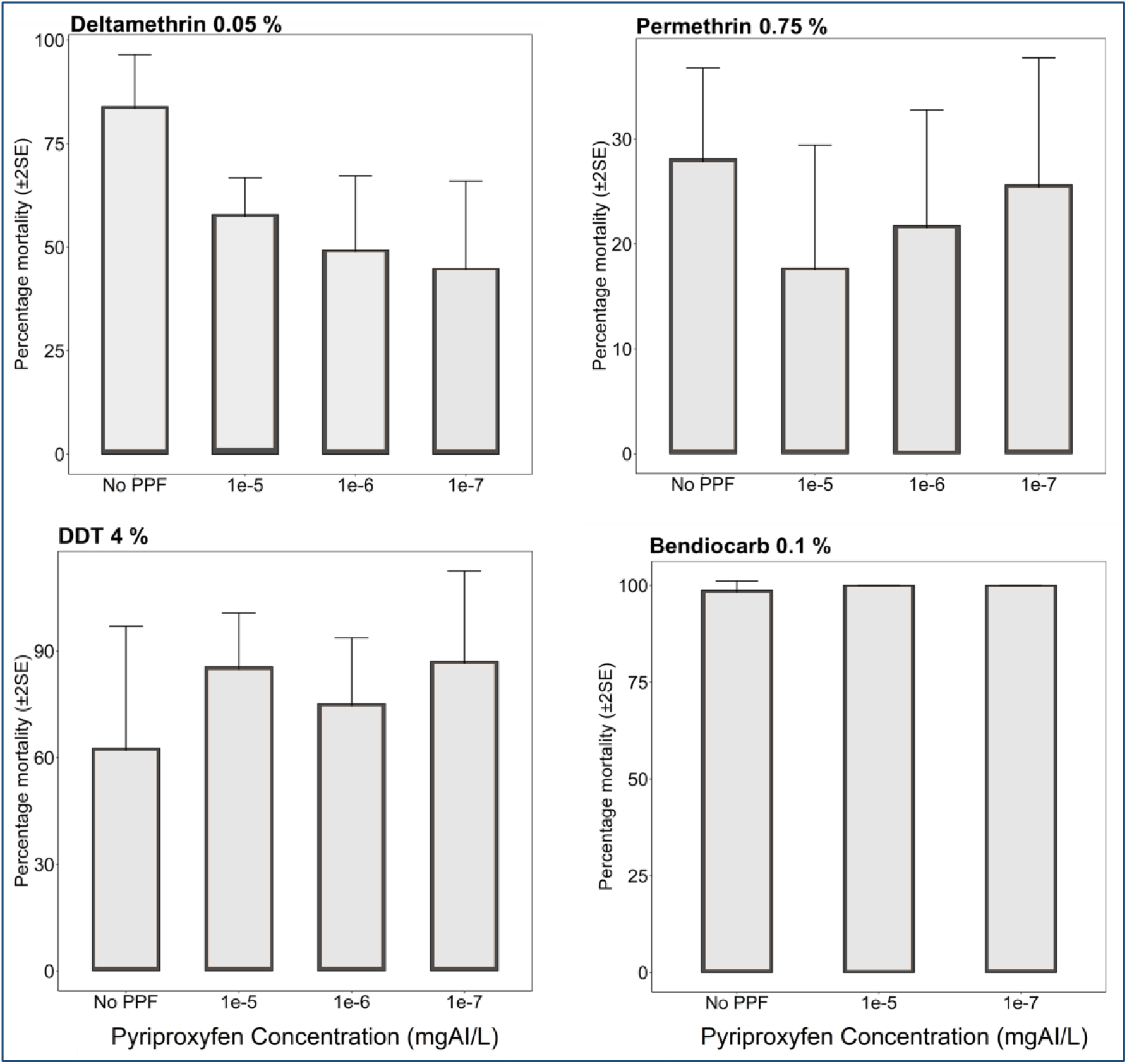
Percentage mortality of wild resistant *An. arabiensis* mosquitoes collected as 3^rd^ instar larvae reared through different doses of PPF (mg of active ingredient per litre, i.e. mgAI/L) and exposed to commonly used insecticides using standard WHO susceptibility testing guidelines. “No PPF” control represents % mortality of wild mosquitoes from the same village, which was not treated with PPF but was exposed to the same insecticide at standard WHO-recommended dose.

We consistently observed reduced 24-hr mortality rates in emergent adults from PPF-treated pools compared to emergent adults from non-treated pools. When 3^rd^ instar larvae of the laboratory-reared pyrethroid-resistant *An. arabiensis* were reared in pools containing different PPF concentrations, phenotypic resistance against permethrin increased, with 24-hr mortality of emergent adults ranging from 5% to 27% (for PPF doses between 1×10^-4^ mg AI/L and 1×10^-9^ mg AI/L) compared to the 35% mortality observed without pre-exposure to PPF in these-laboratory-reared mosquitoes (Fig. 3). Similarly, when the *An. arabiensis* colony emergent adults from different PPF doses exposed to permethrin were pooled, the 24-hr mortality of emergent adults was significantly different (13 %) compared to 35 % observed without pre-exposure to PPF (Fig. 2). Resistance against deltamethrin was also higher, with 24-hr mortality of emergent adults ranging from 26 % to 55 % (for PPF doses between 1×10^-4^ mg AI/L and 1×10^-8^ mg AI/L) compared to 83 % mortality observed when the mosquito larvae had not been exposed to PPF (Fig. 3). On the other hand, when the *An. arabiensis* colony emergent adults from different PPF doses exposed to deltamethrin were pooled, the 24-hr mortality of emergent adults with pre-exposure to PPF (48 %) was significantly different compared to those not pre-exposed to PPF (83 %) (Fig.2).

Similarly, in tests using field-collected mosquitoes, permethrin resistance of *An. arabiensis* emerging from PPF-treated pools was significantly higher compared to that observed in mosquitoes without prior PPF-exposure (Figs. 2 & Fig.4). The 24-hr mortality of these mosquitoes against permethrin ranged from 18 % to 25 % (for PPF doses between 1×10^-4^ mg AI/L and 1×10^-7^ mg AI/L) compared to 28 % without PPF exposure (Fig. 4). We also observed that the 24-hr mortality of the field-collected mosquitoes against deltamethrin ranged from 48 % to 58 % (for PPF doses between 1×10^-4^mg AI/L and 1×10^-8^mg AI/L) compared to 85 % without PPF exposure (Fig. 4). On the other hand, the laboratory-reared *An. arabiensis* remained susceptible to DDT (24-hr mortality = 98%) and bendiocarb (24-hr mortality = 100%), before and after exposure to PPF (Fig. 4). There was also no observable PPF effect when the field-collected mosquitoes were exposed to bendiocarb (24-hr mortality = 100%, compared to 98% before PPF exposure) (Fig. 4). All these observations of increasing resistance levels were made on mosquitoes of the same filial generation, i.e. on mosquitoes whose larvae had been exposed to PPF.

### LLIN bio-efficacy assays

In the WHO cone bioassays, exposure of *An. arabiensis* mosquitoes for 3 minutes to new Olyset^®^ Net resulted in 24-hr mortality of 45%. However, after rearing the same species in larval rearing basins treated with PPF doses ranging from 1×10^-8^ mg AI/L to 1×10^-4^ mg AI/L, there was a reduction in bio-efficacy of the LLIN (Fig.5). The 24-hr mortality of mosquitoes emerging from PPF-treated pools ranged from 15 % to 25 % for Olyset^®^ Net and (30 % to 60 %) (Fig. 5). When the *An. arabiensis* pre-exposed to PPF were pooled for all concentrations and compared with those without pre-exposue to PPF, there was significant reduction in 24-h mortality following exposure to Olyset^®^ Net (t=8.586, df=13.214, p=<0.001). On the other hand there was no significant differences in 24-h mortality when exposed to Permanet (t=0.216, df=5.248, p=0.8365). As in the WHO susceptibility assays, all these observations of reduced bio-efficacy were made on mosquitoes of the same filial generation, i.e. on mosquitoes whose larvae had been exposed to PPF.

**Fig. 5:**
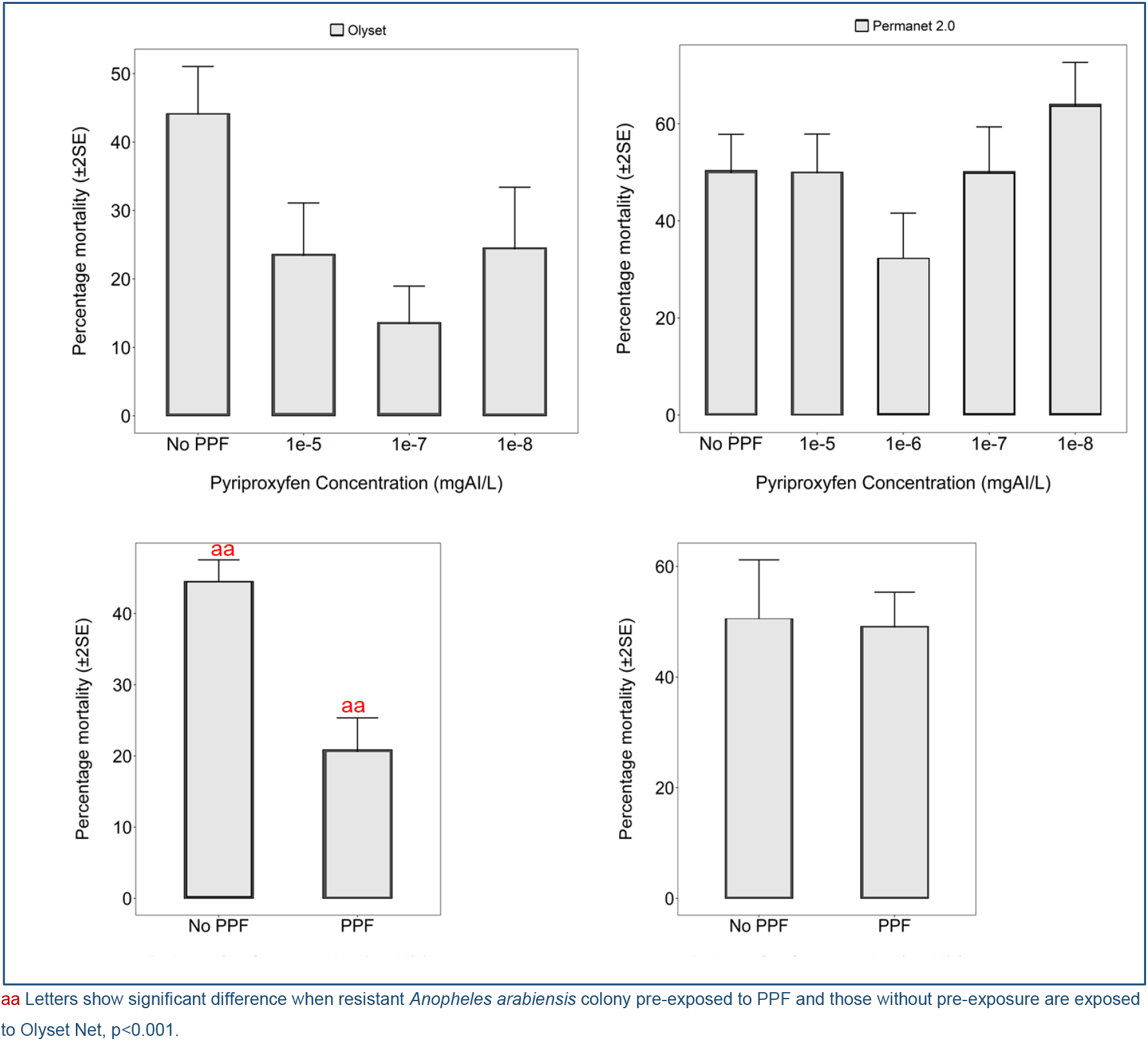
Bio-efficacy (% 24-hr mortality) in cone bioassays of WHO-recommended Olyset^®^ Net against colony-derived pyrethroid-resistant *An. arabiensis* mosquitoes, collected as 3^rd^ instar larvae and reared through different doses of PPF. “No PPF” represents % mortality of mosquitoes from the same colony, which was not treated with PPF but was exposed to the same net.

### Tests using the synergist, Piperonyl Butoxide (PBO), to assess the biochemical basis of resistance

In tests conducted using laboratory-reared *An. arabiensis* mosquitoes, phenotypic resistance to both deltamethrin (24-hr mortality = 58%) and permethrin (24-hr mortality = 23%) was confirmed. Exposure to 4% PBO followed by permethrin or deltamethrin resulted in significantly higher mortalities of the test mosquitoes (24-hr mortality = 85% for permethrin, and 24-hr mortality = 98 % for deltamethrin (Fig. 6). The associated percentage 60-minute cumulative knock-down among the mosquitoes after exposure to the pyrethroids was 96 % for permethrin and 86 % for deltamethrin (Figs. 6 & 7). Comparing tests with PBO exposure to tests without the PBO exposure, the amount of time taken to knock-down 50% of laboratory-reared *An. arabiensis* (KD50) was significantly reduced from 51.10 to 26.86 minutes for deltamethrin and 72.04 to 26.25 minutes for permethrin, the shorter periods having been observed in the tests with PBO exposure (Fig.7).

**Fig. 6:**
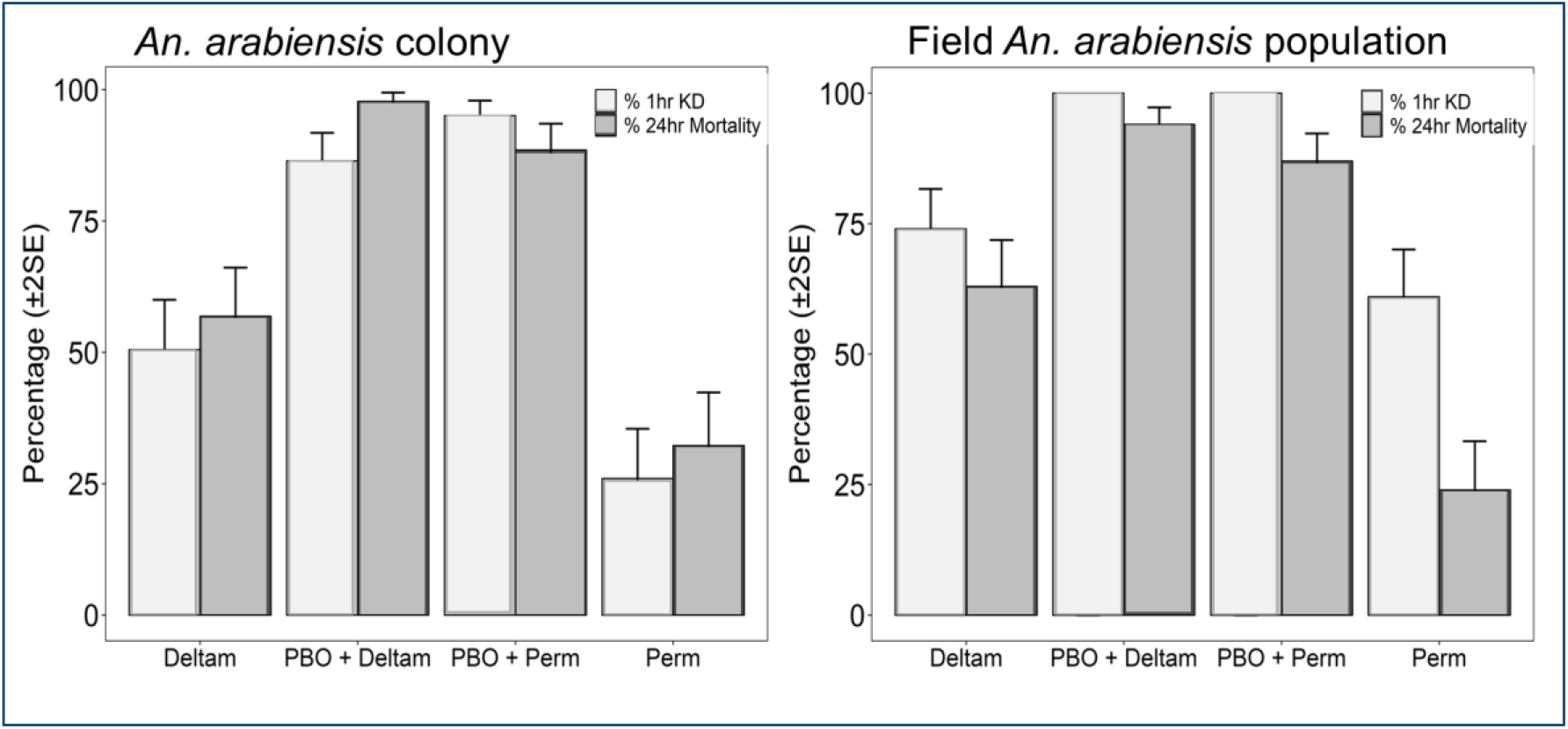
Knockdown time and mortality rates of resistant: *An. arabiensis* colony population and *An. arabiensis* wild population. Increase in mortality and knockdown rate (KD) to 0.05% Deltamethrin and 0.75% Permethrin was observed following 1-hour pre-exposure to synergist; PBO=P450 inhibitor.

**Fig. 7:**
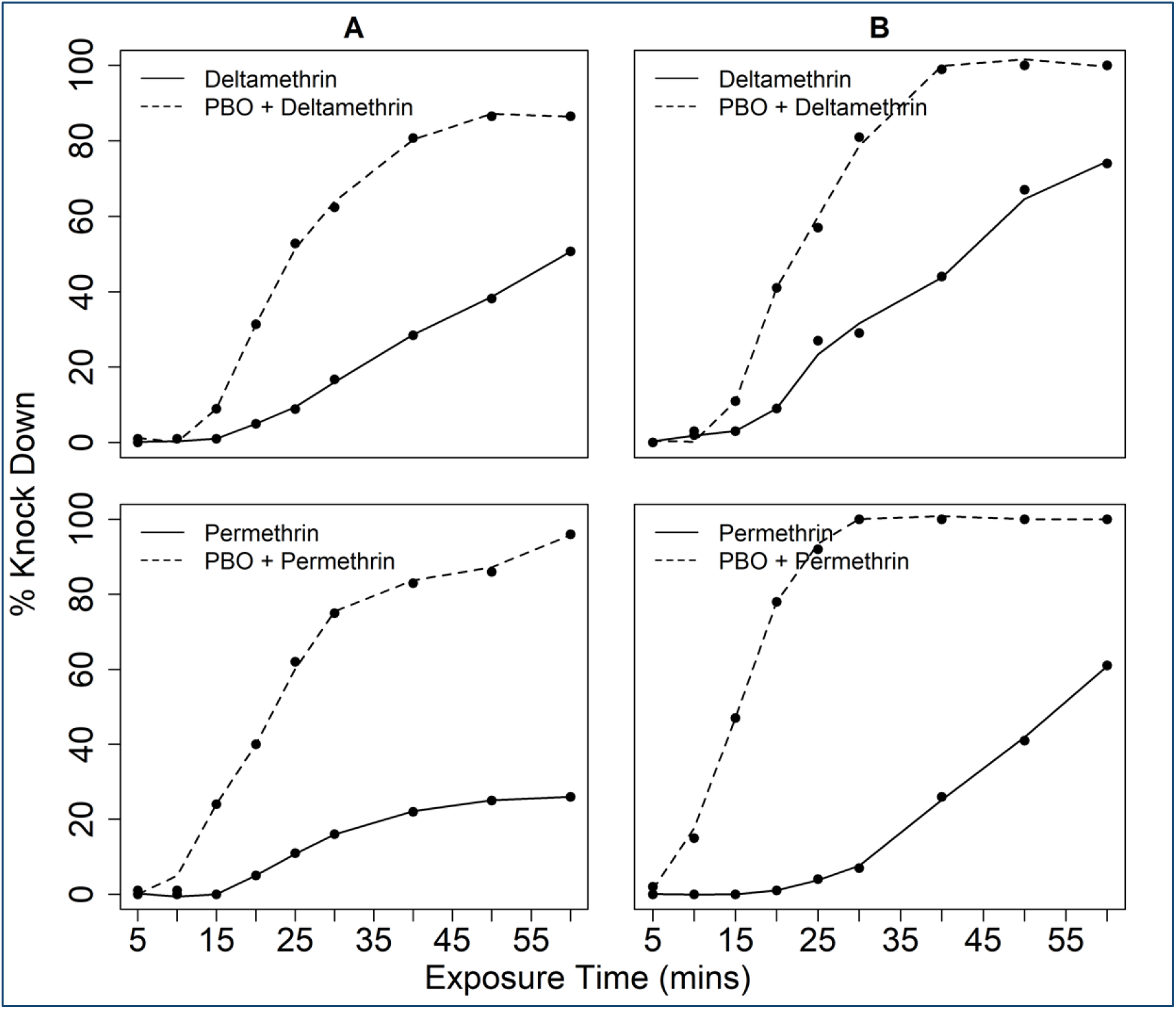
Knockdown time responses of resistant: **A.** *An. arab*iensis colony population and **B.** *An. arabiensis* wild population to **0.05 % Deltamethrin** and **0.75 % Permethrin.**

In tests using field-collected *An. arabiensis* mosquitoes, exposure to deltamethrin and permethrin alone resulted in 24-hr mortalities of 63% and 23% respectively, also confirming the pyrethroid resistance in the study area. However, when these field-collected mosquitoes were first exposed to PBO, then exposed to deltamethrin or permethrin, mortality significantly increased to 95 % and 85 % respectively, accompanied by 100 % cumulative knockdown in each case (Figs. 6 & 7). Similarly, the KD50 for the field-collected *An. arabiensis* population reduced from 43.25 to 23.26 minutes for deltamethrin and permethrin 53.76 to 15.82 minutes, the shorter periods having been observed in the tests with PBO exposure (Fig.7).

## Discussion

The search for new insecticides and insecticide delivery formats to combat the rapid spread of pyrethroid resistance among disease-transmitting mosquitoes is a priority public health agenda. There is growing interest to use juvenile hormone analogues particularly PPF, as larvicide to reduce immature stages of resistant malaria vectors [36, 38]. While several studies have demonstrated the field efficacy of PPF on both the aquatic insects and the emergent adults, little has been done to assess how PPF-based interventions would interact with other vector control measures in the communities. This is particularly important in areas where pyrethroid based intervention, such as LLINs and IRS are already widely used, and where there are possibilities of cross-resistance between PPF and pyrethroids [45]. We, therefore, conducted a study to explore the impact of PPF when added to larval habitats of malaria vectors, and whether that can reduce or increase levels of resistance among these mosquitoes. We examined these interactions in situations where the emergent adults are exposed to different insecticide classes commonly used for public health, i.e. Types I and II pyrethroids, carbamates, and organochlorides. The findings suggest that the addition of sub-lethal doses of PPF to habitats of pyrethroid-resistant larvae, render the emergent adults that evade the aquatic effects, more resistant to pyrethroids (deltamethrin and permethrin) than without such PPF exposure. We observed that this effect is obvious where the mosquitoes are already at least partially resistant. Moreover, we also observed that these negative effects occurred only against pyrethroids but not carbamates (bendiocarb) or organochlorides (DDT). These findings suggest that PPF confers crossresistance to pyrethroids and that aquatic exposure to sub-lethal doses of PPF may exacerbate pyrethroid resistance levels in the malaria mosquitoes. Crossresistance has been previously reported in other insects in studies using selected pyriproxyfen-resistant *Musca domestica* to various insecticide classes, including diacylhydrazine such as methoxyfenozide; triazines such as cyromazine, and benzoylureas such as lufenuron [46, 47]. In another study that explored interactions between PPF and pyrethroids, in the malaria mosquito, *An. arabiensis* found a subset of elevated enzymes responsible for detoxification of pyrethroids that could also metabolize PPF [45], raising concerns that exposure to one may lead to cross-resistance to the other class of compounds. Elsewhere, it was observed that over-expression of one cytochrome P450 gene; *CYP4G61* was associated with resistance to PPF in the greenhouse whitefly, *Trialeurodes vaporariourum* which is a common virus vector and vegetable pest in temperate regions [48].

In our study, we also observed that increased resistance occurred only in cases where the mosquitoes were already showing signs of resistance or reduced susceptibility to pyrethroid. The initial over-expression of the responsible detoxification enzymes may likely have been induced by pyrethroid exposure but that sub-lethal aquatic exposure to PPF only magnified these effects. Previous studies have shown that multiple P450 genes can be concurrently up-regulated in insecticide-resistant mosquitoes through both constitutive transcriptional overexpression, and induction by both exogenous and endogenous compounds, enhancing their role in the detoxification of insecticides in the target organisms [49]. The likely result is that combining PPF with pyrethroids for vector control may create a vicious cycle where resistance to both classes of compounds commonly used is increasingly and rapidly enhanced and magnified. This is also likely to be magnified especially as the insecticide bio-availability wanes overtime. It is important to note that all the observations of increasing resistance levels were made on mosquitoes of the same filial generation. This suggests that elevation of responsible detoxifying enzymes during the aquatic stages can have a significant impact in the mosquito adult stages and that such effects are immediate, and exogenously amplified.

Our study also indicated that mortality rates recorded during the bio-efficacy assessments of the WHO-recommended LLINs Olyset^®^ net was much lower in mosquitoes having prior-aquatic exposure to PPF than mosquitoes without PPF exposure. This suggests having sub-lethal doses of PPF in the environment could compromise the efficacy of these nets. Reduction in the bio-efficacy of standard LLIN against pyrethroid-resistant mosquitoes have been reported in many studies across Africa [50–52], and also in Tanzania [53], near the villages where larvae for this current study were collected. Other than target site mutations, the most common underlying cause of this resistance is the overexpression of cytochrome P450 genes [49]. In contrast to our study, none of the previous studies have evaluated the bio-efficacy of insecticide-treated bed nets, in the context of PPF-pyrethroid use. No previous studies have attempted to investigate the negative cross-resistance effect when PPF is introduced in resistant malaria vectors larval stages. Nevertheless, given these initial results, further studies must be conducted to evaluate the bio-efficacy of other WHO-recommended LLIN including the new generation LLINs against disease-transmitting mosquitoes, and what the overall impact of large scale PPF use may mean for pyrethroid-based vector control. While bed net coverage increases and pyrethroid resistance is documented in nearly all regions endemic to malaria across Africa [54], compounds with effects such as we have observed here with PPF might further compromise the efficacy of these tools.

Despite these worrying findings of how PPF and pyrethroids might interact to magnify resistance and compromise interventions, it is also possible that our study might have underestimated the impact of these interactions. This is because the resistant mosquito larvae were exposed to PPF for only a short duration (starting from 3^rd^ instar larval stage to adult emergence), yet this might not be the case in real field scenarios, where the impact of PPF is likely to build up over time and the mosquitoes are likely to be exposed for longer periods. Therefore, further studies must be conducted to understand the impact of this interaction through selection pressure with PPF over generations with pyrethroid-resistant mosquitoes. In this current study, all observations were made on mosquitoes of the same filial generation, and not after multiple generations of exposure. It is also worth noting that where suppression of mosquito populations resulting from PPF is high enough, it may be adequate to suppress transmission significantly despite the exacerbation of resistance levels. In addition, to minimize the impact of sub-lethal impact against pre-exposed mosquitoes, the use of PPF products may be resolved by timed applications, rather than perpetual applications.

Combining synergists with insecticides used in public and agricultural health sectors, particularly PBO with pyrethroid has long been demonstrated to effectively enhance the efficacy of pyrethroids through inhibiting metabolic enzymes as well as enhancing cuticular penetration in insects [42, 55]. In our study, we also investigated whether metabolic resistance was contributing to both wild and colonized *An. arabiensis* population’s phenotypic resistance. Our results demonstrated that pre-exposure of PBO followed by either deltamethrin or permethrin for both wild and colonized pyrethroid-resistant mosquitoes appeared to reverse the high resistance to these two insecticides, though this reversal was not up to 100% level. Moreover, the cumulative percentage knock-down induced by the two insecticides was recovered to 100% with exposure to PBO first for field-collected mosquitoes and to between 90% (deltamethrin) and 97% (permethrin) for laboratory-reared mosquitoes. The synergism effects of PBO with pyrethroids observed in this study indicate that P450s play a major role in deltamethrin and permethrin resistance in both the laboratory-reared and field-collected *An. arabiensis* populations. Moreover, likely, P450s may also be involved in PPF resistance in the mosquitoes. Taken together with our current study, an increase in mortality and KD following PBO pre-exposure with deltamethrin has been reported in *An. gambiae* s.l and other mosquito vectors elsewhere [42, 56–58]. The reduction in KD time and tolerant levels observed has been associated with increased pyrethroid cuticular penetration in the presence of PBO synergist [42, 43, 55]. It should be noted that in our study the mortality was not recovered to 100% with PBO pre-exposure in both field and colony populations, indicating that alongside metabolic resistance, there could be other resistance mechanisms involved, and further studies are necessary to understand these mechanisms in detail.

World Health Organization (WHO) recommends the use of diagnostic dose assays with fixed exposure times to determine insecticide resistance in a mosquito population. Nevertheless, such discriminatory doses alone cannot be used to determine if changes in the intensity of resistance exist in a population. To quantify further the impact of using PPF on the pyrethroid-resistant mosquito population, we therefore, recommend further studies with varied exposure times of resistant mosquitoes reared or not in PPF to permethrin and deltamethrin. Further studies are also required to assess the delayed effect on mosquitoes beyond 24-h period following pre-exposure to PPF. To the best of our knowledge, no study has assessed changes in resistance to pyrethroids in the context of PPF interaction. The few studies that exist have looked at the intensity of resistance in pyrethroids alone by either varying exposure times or insecticide concentrations and have reported increased strength of resistance in mosquito population to pyrethroids over time [59, 60]. These intensity studies are crucial to inform further the implications of combining PPF and pyrethroid based interventions. It is also important that these further studies on the intensity of resistance are conducted using different methods to link the outcome with the effectiveness of pyrethroid based tools that are currently in place for control of malaria vector mosquitoes. For example, a more realistic picture would be to conduct a semi-field system (SFS) experiment where resistant mosquitoes are released into the SFS in the presence of either sprayed huts with pyrethroids or the presence of LLIN’s at the same time providing artificial breeding habitats treated with sub-lethal doses of PPF and the impact monitored after a series of generations. Another example would be to use tunnel assays where mosquitoes are exposed to a net in the presence of a host and this would provide more information on mortality rate and other biological indicators such as blood-feeding inhibition.

Our findings herein are expected to help guide selections of interventions that combine different interventions at different stages to obtain an optimal integrated vector management approach against mosquito-borne diseases mitigating insecticide resistance. For example, PFF did not onset pyrethroid resistance in otherwise susceptible mosquitoes suggesting PPF could only be used where no pyrethroid resistance has arisen. Alternatively, since we did not observe any negative interactions with bendiocarb or DDT, it might be appropriate to consider integrating PPF-based larviciding with non-pyrethroids for adult mosquito control. One limitation of our study was that we did not assess whether the emergent mosquitoes surviving the sublethal effects of PPF, though more resistant to pyrethroids, were also less sterile or not and for how long they would survive in the wild. PPF is known to function by reducing longevity, the fecundity of mosquitoes, but also by sterilizing the adults. Such sterilizing benefits may remain obvious even if the mosquitoes are themselves more resistant. It would be important therefore to examine the overall impact of combining PPF and pyrethroid-based interventions, considering all modes of action of the active ingredients. Of particular interest would be the long-term effects of next-generation LLINs such as Olyset^®^ Duo, which has both PPF and permethrin as active ingredients against host-seeking adult mosquitoes.

## Conclusions

Exposure of pyrethroid-resistant mosquitoes to sub-lethal aquatic concentrations of PPF increases their levels of resistance. Such effects are apparent in cases where the mosquitoes have any signs of resistance to the pyrethroids. The effects are also not observable against carbamates and organochlorides. Efficacy of pyrethroid-based interventions such as LLINs can be compromised by such sub-lethal exposures. These effects are rapid and occur within a single filial generation, i.e. on adults emerging from contaminated larvae in the same generation. Field applications of PPF, either by larviciding or other means, could potentially magnify levels of pyrethroid-resistance, in areas where signs of such resistance already exist in the vector populations. These findings could help guide the selection of chemicals for integrated control of mosquito larvae and adults, to achieve optimal combinations mitigating the increasing threat of resistance.

## Acknowledgments

We greatly thank Andrew Kafwenji, Paulina Sanga, Nehema Nombo, and Joseph Mgando for continuously providing mosquitoes for our experiment. We also thank Dr. Dickson Lwetoijera of IHI for providing useful advice during the initial stages of this experiment and Mr. Paul Kaefer for his useful comments on this manuscript. This work was funded by the Bill and Melinda Gates Foundation (Grant Numbers: OPP52644). N.S.M received financial support from Wellcome Trust and the Association of Physicians of Great Britain and Ireland (Grant number: WT104029/Z/14/Z), while H.S.N, and F.O.O were supported by a Wellcome Trust Intermediate Research Fellowship (Grant number: WT102350/Z/13/Z) and a grant from World Health Organization, Special Program for Research and Training in Tropical Diseases (Grant No. B40445). We acknowledge the support from the Spanish Ministry of Science and Innovation through the “Centro de Excelencia Severo Ochoa 2019-2023” Program (CEX2018-000806-S), and support from the Generalitat de Catalunya through the CERCA Program. CISM is supported by the Government of Mozambique and the Spanish Agency for International Development (AECID).

## Authors’ contribution

MAO, FOO conceived and designed the study. MAO, SAM, MM, NSM performed the experiments. MAO, HSN performed the statistical analysis. MAO prepared the initial manuscript draft. MAO, FOO critically reviewed the final draft of the manuscript. MAO, HSN, NSM, SM, FOO contributed materials. All authors discussed the results and contributed to the final manuscript.

